# The effect of illumination cues on color constancy in simultaneous identification of illumination and reflectance changes

**DOI:** 10.1101/2024.12.05.626994

**Authors:** Lari S. Virtanen, Maria Olkkonen, Toni P. Saarela

## Abstract

To provide a stable percept of the surface color of objects, the visual system needs to account for variation in illumination chromaticity. This ability is called color constancy. The details of how the visual system disambiguates effects of illumination and reflectance on the light reaching the eye are still unclear. Here we asked how independent illumination and reflectance judgments are of each other, whether color constancy depends on explicitly identifying the illumination chromaticity, and what kinds of contextual cues support this identification. We studied the simultaneous identification of illumination and reflectance changes with realistically rendered, abstract 3D-scenes. Observers were tasked to identify both of these changes between sequentially presented stimuli. The stimuli included a central object whose reflectance could vary, and a background that only varied due to changes in illumination chromaticity. We manipulated the visual cues available in the background: local contrast and specular highlights. We found that identification of illumination and reflectance changes was not independent: While reflectance changes were rarely mis-identified as illumination changes, illumination changes clearly biased reflectance judgments. However, correct identification of reflectance changes was also not fully dependent on correctly identifying the illumination change: Only when there was no illumination change in the stimulus did it lead to better color constancy, that is, correctly identifying the reflectance change. Discriminability of illumination changes did not vary based on available visual of local contrast or specular highlights. Yet discriminability of reflectance changes was improved with local contrast, and to a lesser extent with specular highlights, in the stimulus. We conclude that a failure of color constancy does not depend on a failure to identify illumination changes, but additional visual cues still improve color constancy through better disambiguation of illumination and reflectance changes.

## 1 Introduction

In photopic light levels, a prominent feature of our visual world is color, a psychological correlate for variations in the relative wavelength composition of light. Determining the colors of objects in our visual environment is complicated by the fact that the spectral power distribution of reflected light depends both on the spectral reflectance of the surface and the spectral power distribution of illumination. Furthermore, the illumination in the environment varies substantially with the time of day, weather, and whether the attended location is in direct sunlight or shaded. Thus, to infer the surface colors of objects, the visual system needs to constantly disambiguate illumination from reflectance (von Helmholtz, 1924), and its remarkable ability to perform this function is called *color constancy*.

The basic mechanisms underlying human color perception, such as trichromacy and cone-opponency, are well-known (e.g., Gegenfurtner & Kiper, 2003). Yet, even after decades of research efforts, we do not fully understand how the visual system produces color constancy (for reviews, see: Brainard & Radonjić, 2014; Foster, 2011; Olkkonen & Ekroll, 2016; Witzel & Gegenfurtner, 2018). Early accounts included reliance on chromatic adaptation where the illumination chromaticity sets the new perceptually achromatic point through sensory adaptation (see, e.g., Hering, 1920). The independent effects of sensory adaptation were later found to be much more limited and supplemented by strong influences of stimulus surroundings (Jameson & Hurvich, 1959, 1961, 1964; Wallach, 1948). Chromatic adaptation mechanisms can be seen to operate in both slow and fast timescales (Fairchild & Reniff, 1995) with an instantaneous component that specifically influences color appearance (Rinner & Gegenfurtner, 2000). The instantaneous effect is attributed to color contrast by an object’s immediate background also referred to as *local contrast* (see, e.g., Hurlbert & Wolf, 2004).

Another line of study focuses on different heuristics based on image properties that the visual system could employ in producing color constancy. For example, early computational algorithms used assumptions about image statistics to estimate the illumination chromaticity and to discount its influence to produce surface color estimates (white patch and edge contrasts: Land and McCann, 1971; average color: Buchsbaum, 1980). However, observers show poorer accuracy in estimating illumination compared to surface color, and the two estimates are not well correlated (e.g., Granzier et al., 2009; Rutherford & Brainard, 2002). Other scene statistics have been proposed to help disambiguate changes of illumination and reflectance chromaticity while not providing a direct estimate of illumination chromaticity (mean cone excitation ratios: Foster and Nascimento, 1994; luminance-redness correlation: Golz and MacLeod, 2002). Additionally, the regularities of spectral variation in illumination and surfaces of natural environments can provide heuristics to constrain the problem of color constancy (e.g., Foster, Amano, & Nascimento, 2006; Foster, Amano, Nascimento, & Foster, 2006; Lee, 1986; Maloney & Wandell, 1987; Morimoto et al., 2016, 2021).

Which cues within a visual scene contribute to color constancy remains actively studied (see, e.g., Brainard et al., 2018; Foster, 2011). Adaptation within a smaller area around the stimulus and across the whole visual scene have an influence at different timescales (Smithson & Zaidi, 2004). However, neither local adaptation, global adaptation, nor adaptation to the brightest spot can fully account for observers’ color constancy (Kraft & Brainard, 1999). Instead, slower sensory adaptation may support adjustment to the average illumination chromaticity in the environment, while faster, regional mechanisms compensate for illumination differences locally (Werner, 2014). Local contrast between adjacent areas has an especially robust effect on how color (and lightness) is perceived (Helson, 1943; Shevell, 1978; Wallach, 1948). Nevertheless, providing an inaccurate illumination cue in the immediate background reduces color constancy but does not eliminate it, pointing to a role for other cues (Delahunt & Brainard, 2004; Kraft et al., 2002). Other stimulus properties, such as three-dimensionality of stimuli, show smaller and less consistent effects (see Mizokami, 2019, for a review). For example, Xiao et al. (2012) found color constancy to be better for flat disks than 3D spheres with consistent illumination cues in the image, but the situation reversed with an inconsistent cue in the immediate background, showing better constancy for 3D spheres. Surface properties, such as glossiness, may also support color constancy by providing information about illumination chromaticity as suggested by computational models (D’Zmura & Lennie, 1986; Lee, 1986; Lehmann & Palm, 2001). Experimental results show that this may be the case, at least when cues from local surround are weak or missing (Granzier et al., 2014; Lee & Smithson, 2016; Wedge-Roberts et al., 2020).

A common method for studying color constancy is asymmetric matching, in which the accuracy of matching surface colors across different illuminations is measured (Arend & Reeves, 1986; Brainard et al., 1997; Burnham et al., 1957). Operational color constancy (Craven & Foster, 1992), on the other hand, refers to the ability to discriminate between changes in illumination and surface reflectance (see, e.g., Foster, Amano, & Nascimento, 2001; Foster, Nascimento, et al., 2001; Linnell & Foster, 1996). Recently, Ennis and Doerschner (2019) studied how observers perceive and estimate simultaneous changes in illumination and object reflectance when shown both an object and a background or either one alone, and how much these estimates rely on globally and locally calculated image statistics. They found that observers are able to resolve both illumination and reflectance when shown the whole stimulus scene but not from the object or background alone. Of the statistics Ennis and Doerschner tested, luminance-redness correlation, mean cone excitation ratios, white patch, and average color could each account for some of the variance in observer responses for either reflectance or illumination. None of the statistics alone was a good general predictor. The authors concluded that the visual system can retain explicit information of illumination changes when a background is present, and that observers are likely to use various sources of information to achieve color constancy. Their experiment did not, however, probe the interdependence of illumination and surface reflectance estimation. Furthermore, the scene surrounding the object was only manipulated between a highly diagnostic white background and no background at all.

In this study, we examine the possible interdependence of illumination and reflectance judgments in operational color constancy. Specifically, even if explicit estimates of illumination chromaticity are inaccurate as previous studies suggest (Granzier & Valsecchi, 2014; Granzier et al., 2009; Khang & Zaidi, 2004), does correct identification of a surface reflectance change depend on explicitly identifying the change in illumination, and if this in turn depends on available visual cues (local contrast or specular highlights). Observers performed simultaneous identification of illumination and reflectance changes with sequentially presented, realistically rendered stimuli. We manipulated available cues for color constancy from local contrast and specular highlights. We found that reflectance judgments were biased by illumination changes. The identification of reflectance changes did not fully depend on explicit identification of illumination changes however, only showing a clear difference in discriminability when there was no illumination change in the stimulus. Identification of reflectance changes was poorer without local contrast cues, and only improved slightly with specular highlights. Identification was also strongly dependent on whether reflectance or illumination chromaticity changed toward or away from neutral.

## 2 Methods

### 2.1 Observers

Three observers took part in a pilot study, and ten new observers took part in the main study. Of the ten observers, nine completed all measurements and were included in data analysis (7 female and 2 male, aged 19-31 with median age of 20). All observers reported having normal or corrected-to-normal visual acuity. Color vision was assessed to be normal with Ishihara color plates (Ishihara, 1987). Prior to the experiment, all observers were given a brief explanation about the interaction of surface reflectance and illumination in producing the light reflected to the eyes. Observers were also told that we were interested in the identification of chromatic surface reflectance and illumination changes in the scene, but they were otherwise naïve to the goals of the study. The study followed the Declaration of Helsinki and was approved by the University of Helsinki Ethical Review Board in Humanities and Social and Behavioural Sciences.

### 2.2 Apparatus

The experiments were carried out in a darkened laboratory room at the University of Helsinki. The experiments were run with custom code on an HP Z230 Desktop PC using Matlab (version R2016b, build 9.1.0.441655) with the PsychToolBox-3 extensions (Brainard, 1997; Kleiner et al., 2007; Pelli, 1997). The display was a 23-inch ViewPixx 3D Lite, controlled by an Nvidia Quadro K620 graphics card. The spectra and gamma functions of the display primaries were measured using a SpectraScan PR-655 spectroradiometer for calibrated color reproduction. Display luminance was linearized by interpolating measured gamma functions. Display white point was set to be metameric to D65, and stimuli were displayed using 100 Hz refresh rate, and 10 bits per primary channel. The high bit-depth display mode that was used for increased color resolution halved the number of horizontal pixels of the native 1920 x 1080 resolution, making the effective display resolution 960 x 1080 pixels. Viewing distance was held constant at 100 cm using a chin rest, and observers gave their responses with a gamepad.

### 2.3 Stimuli

Stimuli were simple, abstract, 3D-rendered scenes (see Figure 1). The objects in the scenes were created by adding irregular Gaussian perturbations to a sphere using ShapeToolbox (Saarela, 2018), and will be referred to as “blobs”. All stimuli included a blob in the middle of the screen, extending approximately 2° in visual angle on average, and we will refer to this central blob as the “target”. Two types of stimuli comprised only a target object with an otherwise black screen (referred to as “object stimuli”, Figure 1a): 1) target flattened in the direction of viewing, “flat object”; 2) matte 3D target, “3D object”. Three types of stimuli comprised a target with different surroundings (referred to as “scene stimuli”, Figure 1b): 1) flat target with a flat gray background, “flat scene”; 2) matte 3D target with four surrounding colored matte objects, “3D matte scene”; 3) matte 3D target with four surrounding colored glossy objects, “3D glossy scene”.

**Figure 1:**
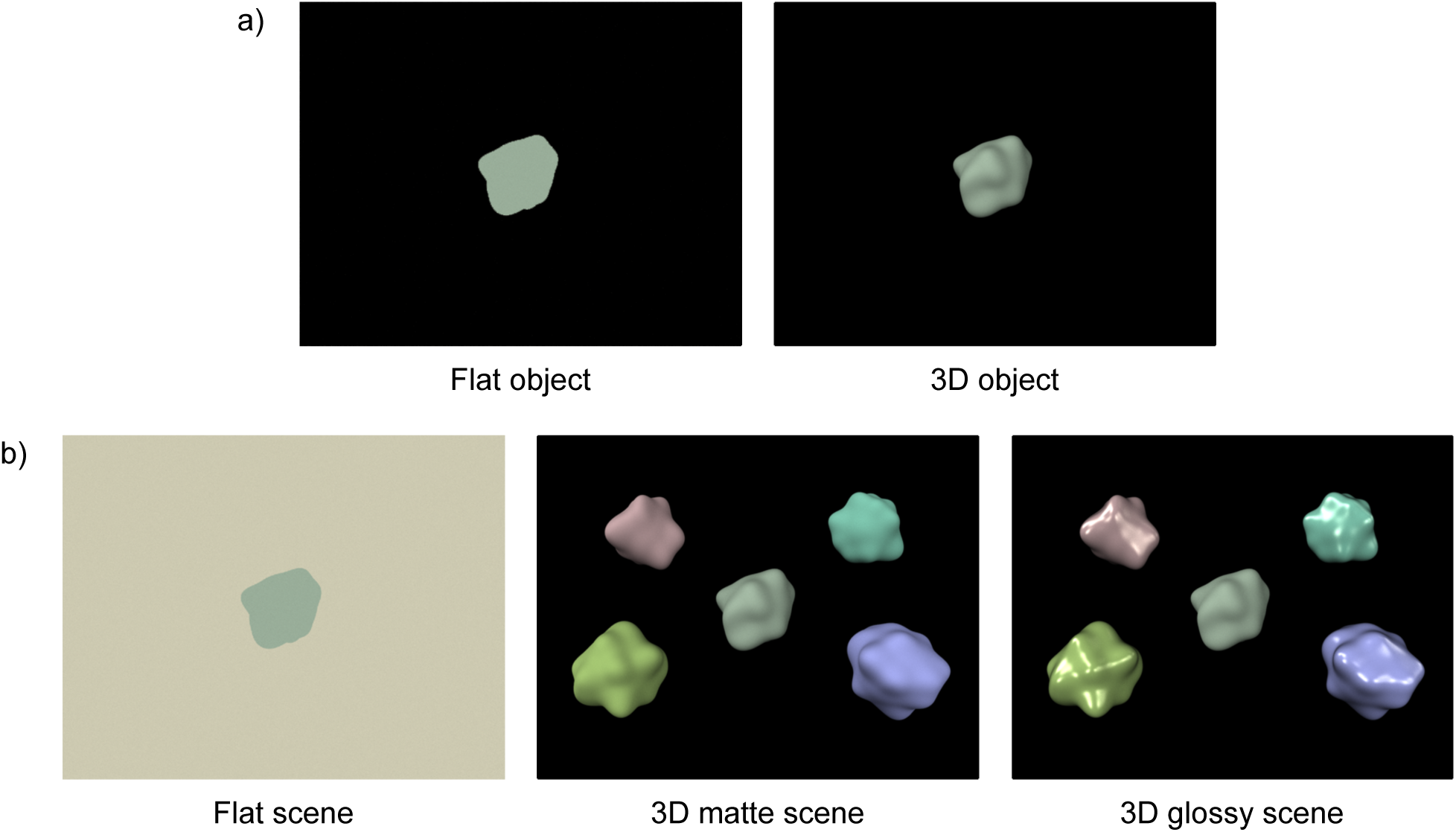
Illustrations of the five different types of stimuli used in the experiment. a) Object stimuli only include a target with an otherwise black screen. b) Scene stimuli include a target and a background affected by changing scene illumination, either as a gray flat surface or additional, colored blobs.

The stimulus scenes were created with spectral, physically based rendering with plain path tracing in Mitsuba (Jakob, 2010) using RenderToolbox3 (Heasly et al., 2014). In all scenes except the 3D glossy scene, all surfaces were modeled with a bidirectional scattering distribution function of an ideally diffuse (Lambertian) material with no specular reflections. In the 3D glossy scene, the target was an ideally diffuse material, while the colored blobs surrounding the target were defined as a plastic-like material with specular and diffuse components (Mitsuba’s default “rough plastic” material). The material modeled a diffuse surface under a thin dielectric layer using Fresnel reflection and transmission coefficients with an interior index of refraction of 1.49 (matching polypropylene). Additionally, the material produced surface roughness by modeling microfacet normals as a Beckmann distribution with root mean square slope of 0.1. Illumination with spatial variation in intensity was created using a freely available high dynamic range light probe as an environment map (“Overcast day at Techgate Donaucity” by Bernhard Vogl, url: http://dativ.at/lightprobes/). The luminance component of the environment map was kept intact while the chromatic component was replaced with the desired uniform chromaticity.

For rendering, we defined discrete spectral distributions between 300 and 700 nm in 10 nm steps for illuminant radiance and the diffuse reflectance component of surfaces (which we will later simply refer to as “reflectance”). First, however, we defined chromaticities for different illuminants and reflectances in CIELUV color space using the *u*^′^*v*^′^-coordinates (the *u*^′^*v*^′^-coordinates for spectral reflectance were defined as the the coordinates of the reflected light when a surface having that spectral reflectance is illuminated by an equal-energy white illuminant). As a measure of chromatic difference between colors we used Euclidian distance in *u*^′^*v*^′^ coordinates (Δ*u*^′^*v*^′^). Note that although CIELUV is only approximately uniform (see, e.g., Mahy et al., 1994; Pointer, 1981), we deemed that equal color differences in Δ*u*^′^*v*^′^ were sufficiently accurate for the purposes of the experiment. D65 chromaticity was always chosen as the “middle” illuminant, and six chromaticities with equal steps in Δ*u*^′^*v*^′^ were chosen along the daylight locus (D-illuminant) in both the lower color temperature (yellower) and the higher color temperature (bluer) directions (Figure 2). For reflectance, 13 chromaticities were chosen so that steps in illumination and reflectance chromaticities matched in the resulting reflected light chromaticity (details below). To create the spectral distributions from *u*^′^*v*^′^ chromaticity coordinates, we used spectral upscaling from XYZ values with basis vectors. For illuminants, we used the CIE D-illuminant components (CIE, 2018). For reflectances we created custom basis vectors by taking the first five principal components from the spectral measurements of Munsell chips by Parkkinen et al. (1989).

**Figure 2:**
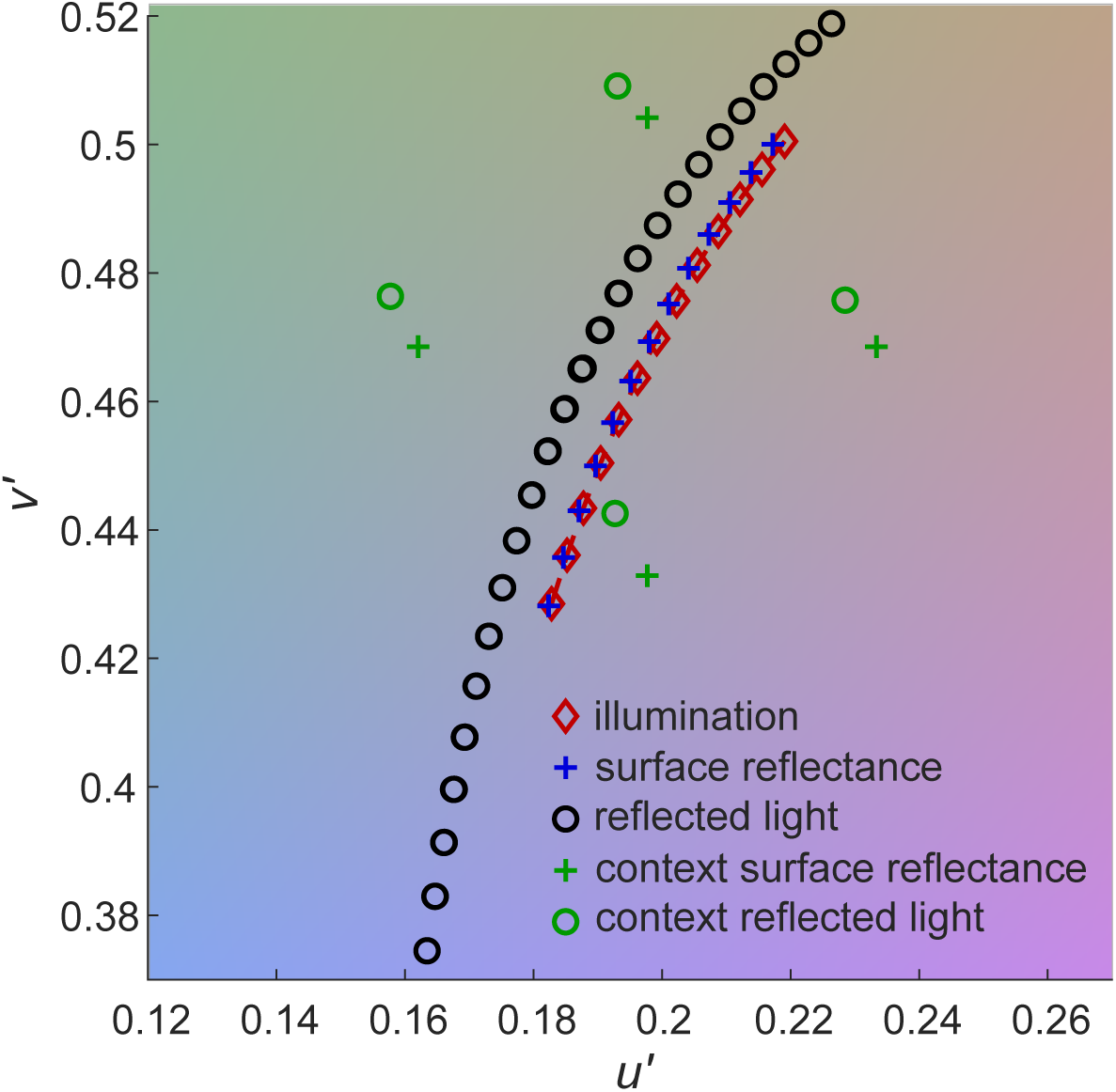
Chromaticities of one of the seven stimulus sets used in stimulus rendering displayed on CIE *u′v′* chromaticity diagram. The *u′*-dimension is on the x-axis, and *v′*-dimension on the y-axis. Red diamonds, blue crosses, and black circles show the 13 different illuminants, 13 different reflectances, and reflected light from all their combinations, respectively (note that multiple circles overlap). Green crosses show the reflectances of the four "context" blobs and green circles their reflected lights with the middle illuminant (D65).

We wanted the reflected target color change to be the same for both illumination and reflectance changes. This was done in order to force reflectance estimation to depend on accounting for possible illumination changes, so that target color alone cannot be used to disentangle illumination and reflectance changes. We chose illuminant and reflectance chromaticities so that a given number of steps leads to the same reflected target color, whether they are illumination or reflectance steps, or both (see Figure 4). Conversely, this also means that opposite changes in illumination and reflectance lead to no change in reflected color. Note that this would not be possible with specular highlights because they reflect the chromaticity of the illumination, which is why we chose to use matte targets in all experimental conditions. To create the 3D glossy scene, we used specular background objects instead.

To estimate suitable chromaticities, we created a custom optimization procedure to minimize errors using Matlab’s fminsearch-function. First, we calculated 13 chromaticity coordinates for reflected light using a reflectance equal to D65 chromaticity with each of the 13 illuminants. We calculated the reflected light by using the spectral upscaling detailed above and element-wise multiplication of illuminant and reflectance spectral distributions. Using reflectances with chromaticity coordinates equal to those of our 13 illuminants as starting points, we searched for reflectance *u*′*v*′ coordinates that would minimize the difference in *u*′*v*′ coordinates of corresponding reflected lights compared to the 13 previously calculated reflected lights, as well as Δ*u*′*v*′ between all adjacent reflectances. Second, we let the *u*′, *v*′, and *Y* (luminance) values of all 13 illuminants and all 13 reflectances vary. We searched for chromaticity coordinates that would minimize the difference between all corresponding reflected lights, the difference in illuminant step sizes (Δ*u*′*v*′), the difference in reflectance step sizes (Δ*u*′*v*′), as well as difference in Y between different illuminants, and between different reflectances. An example of the resulting chromaticity coordinates is illustrated in Figure 2.

Eight sets of such coordinates were created, each with a different step size (approximately 0.0039, 0.0055, 0.0064, 0.0073, 0.0085, 0.0096, 0.0109, and 0.0123 in Δ*u*′*v*′). The set with the smallest step size was only used for the preliminary measurement while the other seven were options for the rest of the experiment (chosen based on the observer’s performance). Finally, chromaticities for the four background blobs were selected by picking coordinates with equal distance as the furthest reflectance steps from D65 in −*u*′, +*u*′, −*v*′, and +*v*′, respectively. For the gray, flat background, reflectance spectrum was always uniform with 0.33 reflectance.

For stimuli without a background, scenes were rendered with the D65 illuminant for all different reflectances, 13 different stimuli for each stimulus set. For stimuli with a background, scenes were rendered with all combinations of illuminant and reflectance spectra, 169 different stimuli for each stimulus set. Additionally, with four surrounding blobs as the background, the blobs were rendered in six different rotations. This was done to discourage observers from consistently using specific areas of the blobs with specular reflections to determine illumination chromaticity, which could have biased spatial attention in the 3D glossy condition.

### 2.4 Procedure

The experiment was divided into three sessions held on different days. The first session included a preliminary discrimination measurement, demonstration (henceforth demo) and practice trials, and one run of the main experiment. Sessions 2 and 3 included a demo and practice set, followed by two runs of the main experiment. We first describe the tasks, and then explain in more detail the content of the sessions.

The aim of the preliminary measurement was to determine a discrimination threshold for stimulus color differences. Observers were shown two flat target stimuli sequentially with no background in 500 ms intervals, separated by a 250 ms inter-stimulus-interval (ISI). The observer responded whether the stimulus in the second interval was yellower or bluer than in the first interval, and received feedback for their response. One of the two stimuli had the median chromaticity, and the other stimulus had one of the 13 chromaticities in the stimulus set. Each chromaticity pair was repeated 13 times, and a cumulative Gaussian psychometric function was fit to the proportion of responses indicating the varying stimulus as bluer. The stimulus set with Δ*u*′*v*′ closest to 1.5 times the observer’s psychometric function standard deviation was chosen for the first practice set.

The main experiment consisted of the stimulus conditions presented in Figure 1, with each condition run in a separate block of trials. Each trial consisted of the presentation of a reference stimulus for 1000 ms, and a comparison stimulus for 1000 ms, with a 250 ms ISI in between (Figure 3). The observer’s task was then to respond how the stimulus colors changed from the reference to the comparison stimulus by moving visual response indicators to their choices with the gamepad’s control sticks and confirming with a button press. In object blocks (Figure 1a), observers responded using a single indicator whether the target surface color had a large change towards yellow, small change towards yellow, no change, small change towards blue, or large change towards blue. In scene blocks (Figure 1b), observers judged both target surface color and scene illumination and responded using separate visual indicators if there was a change towards yellow, no change, or a change towards blue. See Figure 3 for an illustration of the screen with two visual indicators. The order of the indicators (left/right), as well as whether blue or yellow responses were up or down, was alternated between observers. Response time was not limited. After a response, observers received feedback with colored response indicators displayed for 1000 ms in practice and dummy trials but not in main experiment trials. The experiment proceeded to the next trial after a 500 ms inter-trial-interval (ITI).

**Figure 3:**
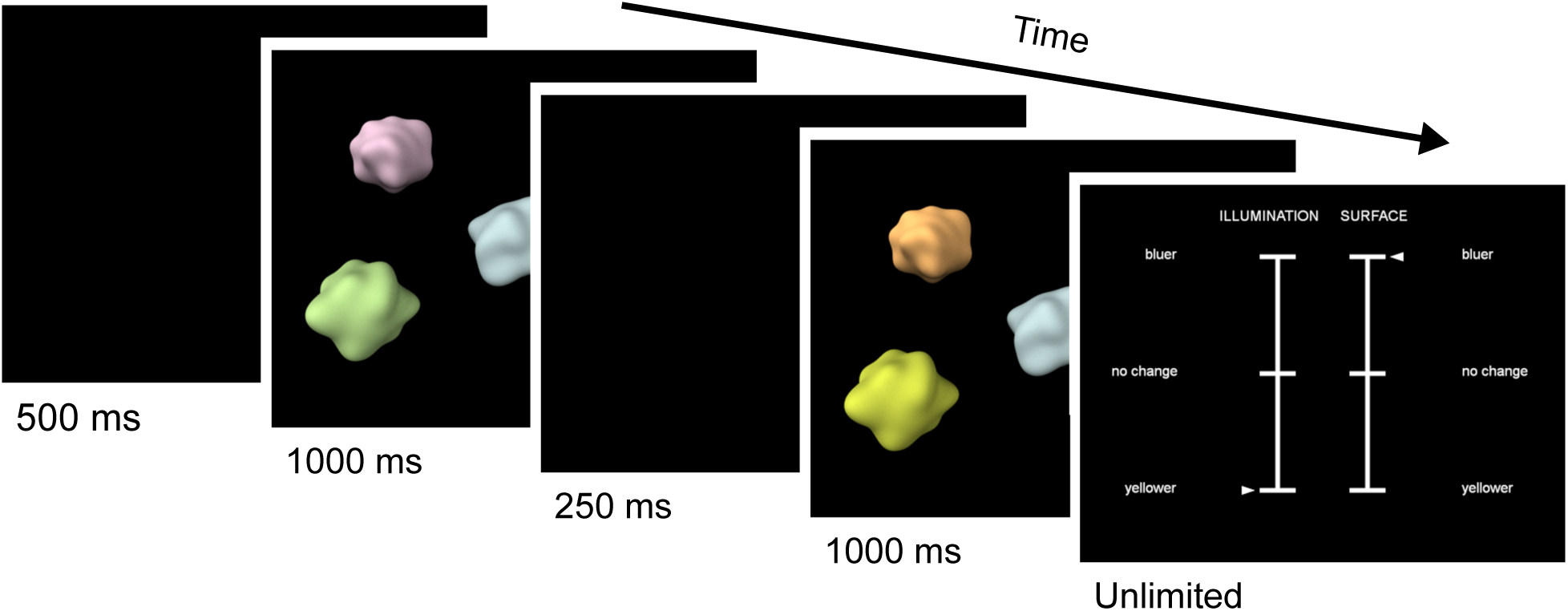
The course of a single trial. The reference stimulus was displayed for 1000 ms, followed by a 250 ms ISI, and the comparison stimulus for 1000 ms. The observer was then shown one or two indicators with written responses alternatives. When the observer gave their response, the experiment moved to the next trial after a 500 ms ITI.

In object blocks, the reference stimulus was chosen from the middle 9 of the 13 reflectance chromaticities (9 different references in total), and the comparison stimulus -2, -1, 0, +1, or +2 chromaticity steps from reference (5 different changes in total). In scene blocks, both illuminant and target reflectance chromaticities were chosen from the middle 9 of the 13 respective chromaticities (81 different references in total), and the comparison stimulus illuminant and reflectance -2, 0, or +2 chromaticity steps from reference (9 different changes in total, see Figure 4).

**Figure 4:**
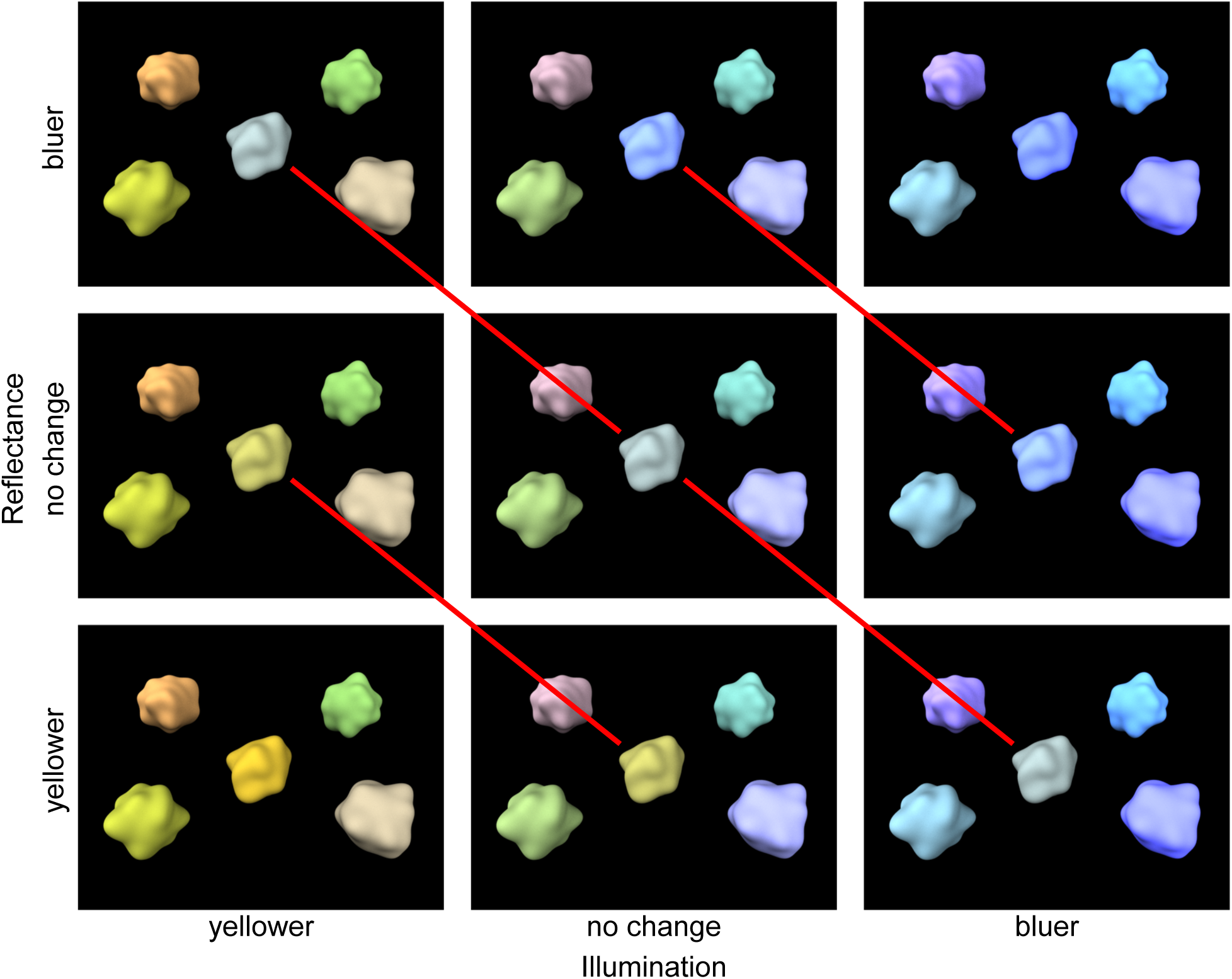
All possible changes in stimulus colors between the reference and the comparison stimulus in scene blocks. In this example, the reference stimulus would be identical to the stimulus in the middle. The comparison stimulus could be one of the nine possibilities in the figure. Changes in illumination are on different columns, and changes in target reflectance are on different rows. The red lines illustrate that similar changes in the illumination or reflectance result in identical changes in the reflected light from the target.

After the preliminary measurement at the beginning of the first session, the rest of the first session and the following two sessions were used to collect data for the main experiment. Each of the three sessions followed the same pattern: first demo trials followed by practice trials for each block in a fixed order, then main experiment trials once for each block in a random order (see Figure 5a). In the second and third sessions main experiment trials were repeated a second time for each block in a random order. In the first session, average *d*′ (*d*′ or marginal *d*′, see below) across all blocks and stimulus changes was calculated from practice trials, and this was used to select the stimulus set for the rest of the experiment. Average *d*′ between 1 and 1.5 led to no change in stimulus set; *d*′ below 1, 0.75, and 0.5 led to a stimulus set 1, 2, and 3 steps higher, respectively; *d*′ above 1.5, 2, and 2.5 led to a stimulus set 1, 2, and 3 steps lower, respectively.

**Figure 5:**
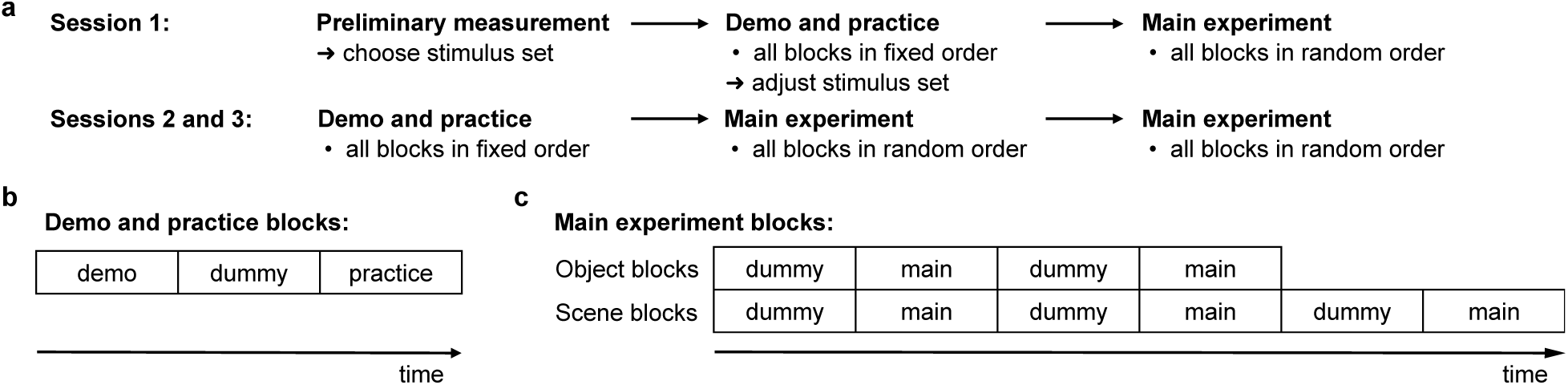
Experiment protocol. a) Measurement session 1 included a preliminary measurement used to choose an initial stimulus set. The following practice trials were analyzed to adjust the stimulus set towards a target level of performance. In sessions 2 and 3, all main experiment trial blocks were repeated twice. b) In demo and practice blocks the practice trials were preceded by demo trials and dummy trials. c) In main experiment blocks, sets of main experiment trials (without feedback) were interleaved with sets of dummy trials (with feedback).

Blocks of demo and practice trials (see Figure 5b) began with the demo trials where observers were first told in writing what kind of stimulus color changes would be demonstrated and were then shown two examples of such trials with different reference stimuli. This was repeated for all possible changes (object blocks) or combinations of changes (scene blocks) in a preset order. The first session had considerably more practice trials than the second and third sessions. In the first session, practice object blocks included 10 trials for each change (50 trials in total). Practice scene blocks included 15 trials for each illumination and reflectance change, shuffled into random combinations (45 trials in total). In the second and third sessions, practice included only two trials for each comparison in object blocks, and 3 trials for each comparison in scene blocks. At the beginning of each practice block, three dummy trials selected randomly from the list of trials were presented.

Main experiment blocks (see Figure 5c) began with and were interleaved by dummy trials with feedback. Object blocks included two dummy sets with one repetition of each change. Scene blocks included three dummy sets with two repetitions for each illumination and reflectance change, shuffled into random combinations. In total there were five repetitions for each main experiment block. Thus each change (object blocks) or combination of changes (scene blocks) had 40 main experiment trials used in analysis.

### 2.5 Data analysis and modeling

First, all dummy trials were discarded from analysis. For overall comparisons between blocks, data were pooled across all references. Based on signal detection theory (Green & Swets, 1966), the object blocks were considered a one-dimensional five-alternative identification task, and scene blocks were considered a two-dimensional three-alternative identification task. Average of *d*′ (or average marginal *d*′ in scene blocks) over change levels was used as a measure of discriminability. Paired *t*-tests were used to compare discrimination between different reference-comparison changes within blocks, and to compare the same reference-comparison changes between blocks. All *p*-values of all statistical tests mentioned were Holm-corrected (Holm, 1979).

To investigate the relationship between identification of illumination and reflectance changes, we pooled the data across observers and stimulus changes within scene blocks and performed chi-square tests of independence for correct illumination and reflectance responses. Additionally, we pooled the data for each scene stimulus across observers only and calculated average marginal *d*′ to identify reflectance changes separately for trials where illumination change was identified correctly and incorrectly. Their equality was tested with significance tests for one-parameter signal detection hypotheses (Gourevitch & Galanter, 1967; Marascuilo, 1970).

In order to explore if identifying changes towards and away from the “neutral” differ, we again used the data pooled across observers. Trials from each of the three scene blocks were binned according to whether the reference stimulus had illumination or reflectance from the yellowest, bluest, or neutral third in the stimulus pool. Within the six trial groups, criterion (*c*) and discriminability (*d*′) were calculated separately for changes towards yellow or blue, and their equality was tested with the aforementioned one-parameter tests (Gourevitch & Galanter, 1967; Marascuilo, 1970).

Finally, we quantified the relationship between identification of illumination and reflectance changes by fitting a two-dimensional signal-detection model to the data from the scene blocks. The model observer makes its decision by comparing noise-corrupted responses to response criteria in a two-dimensional decision space, where the response to a stimulus is given by two hypothetical mechanisms: a “reflectance mechanism”, and an “illumination mechanism”. This model is not intended to be a model of actual mechanisms of color constancy, rather it is used as an analysis tool to quantify the amount of cross-talk in the processing of the reflectance and illumination cues.

We represent the stimulus by a vector ***s*** = [*s*_r_ *s*_i_]^⊤^, where the reflectance value *s*_r_ is negative for a yellow shift, zero for no change, and positive for a blue shift in reflectance. The same holds for illumination changes for the illumination value *s*_i_. The sensitivity of the two mechanisms to illuminance and reflectance changes is given by the vectors ***m***_r_ and ***m***_i_, put in the rows of matrix *M*:

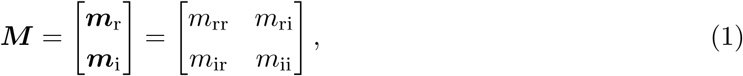

where the first subindex is for the mechanism and the second for the stimulus feature (so that, for example, *m*_ri_ gives the sensitivity of the reflectance mechanism to illumination changes). If illumination and reflectance cues in our stimuli were processed independently of each other, matrix *M* would be diagonal. The noisy responses of the mechanisms are then computed as

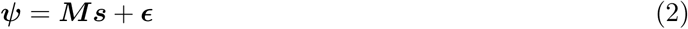

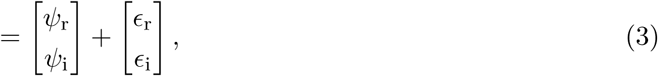

where the noise ***ε*** is zero-mean and normally distributed:

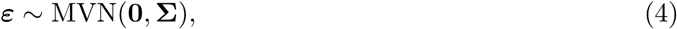

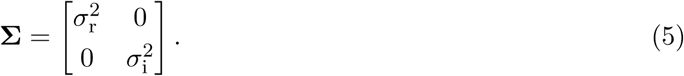

We fixed the noise covariance to zero, because the possible cross-talk between the two stimulus dimensions is accounted for by the mechanism sensitivities (in matrix ***M*** ). We fixed the noise variances to 1 for zero response and assumed that the variance increases linearly with the mechanism response (*σ*^2^ = 1 + *kψ*, where *k* is a constant, for both responses). The model observer makes the final identification decision by comparing the noisy response to two response criteria along each stimulus dimension.

We thus fit ten parameters in total: two parameters to scale the variances, four parameters for decision criteria, and four parameters for the response mechanism sensitivities. Parameters were fit by maximum likelihood (in practice by minimizing the negative log-likelihood of the parameters given the observed responses). Judged by the Bayesian information criterion, this model gave a better fit than a full model where the variances and mean responses are free parameters.

The model allows for perceptual independence to fail between the two stimulus dimensions (because the two mechanisms are not necessarily orthogonal), but it assumes decisional separability (because the decision criteria are parallel to the axes of the decision space). Note that the effect of non-orthogonality of the mechanisms could also be modeled as having orthogonal dimensions but correlated noise (Ashby & Townsend, 1986). We chose to have potentially non-orthogonal mechanisms to better illustrate the dependencies in the processing of the two cues.

### 2.6 Results

Object block response counts summed across observers (Figure 6a–b) show that the correct option had the most responses in all cases except for the large change towards yellow for the flat object. Nearly all of the incorrect responses were in the response options adjacent to the correct option. There was some tendency to report less of a change in the yellow direction than what is shown in the stimulus, both in general and in comparison to changes towards blue. On average, discriminability of color changes in the stimuli (Figure 6c) did not show a statistically significant difference in paired *t*-tests between the two object blocks, or between yellower and bluer changes within blocks.

**Figure 6:**
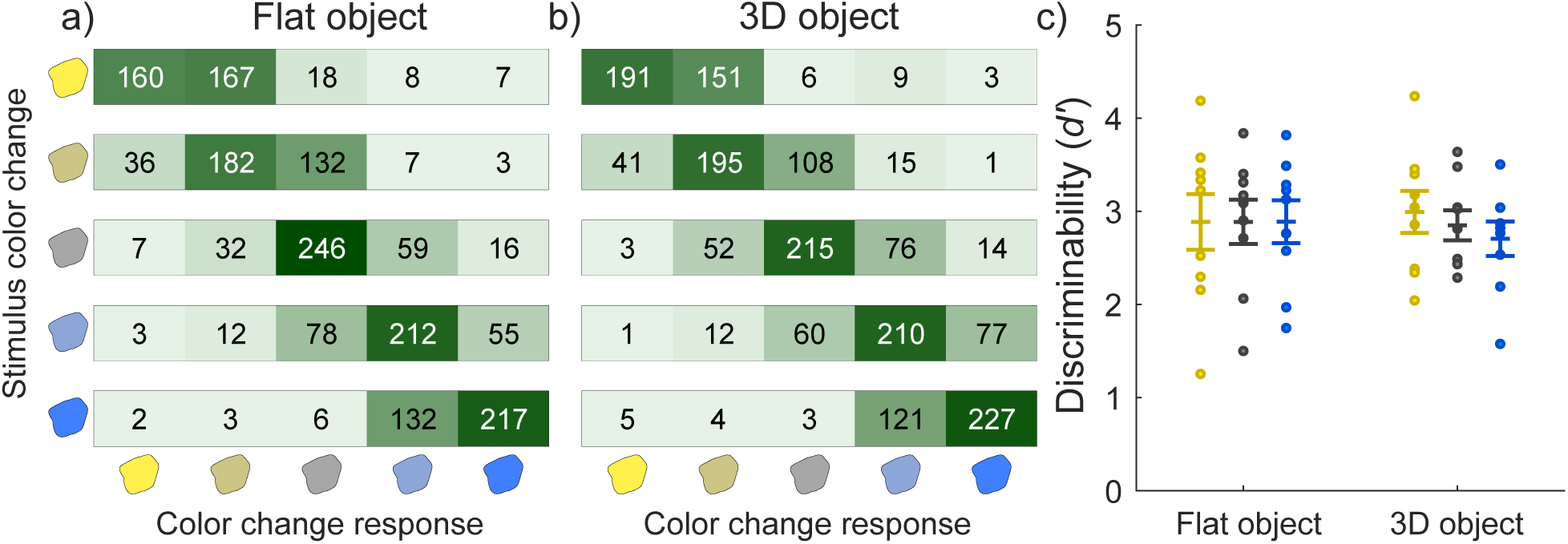
Change identification results from object blocks. Heatmaps of response counts across all observers for: a) flat object, b) 3D object. Each stimulus change is shown on a separate row. Left-hand icons indicate the correct color change response, and bottom icons indicate observers’ color change responses. c) plots discriminability measured by average *d′* between the neutral and the “large” color differences. Different object blocks are on the x-axis with *d′* on the y-axis. Each dot is a single observer, and the error bar indicates ±1 SEM. Yellow dots show averages across changes towards yellow, blue dots towards blue, and gray the total average.

Figure 7a–c shows the response counts from the scene blocks for each of the 3x3=9 possible stimulus changes. The correct option again had the most responses in most cases and incorrect responses were mainly for response options adjacent to the correct option. There was, however, also interaction between the responses to the two types of stimulus change. Most notably, observers were most accurate when there was either no change or a simultaneous change in both illumination and reflectance towards the same color. In contrast, observers performed worst when both illumination and reflectance changed towards opposite colors (and the reflected color thus stayed the same), often reporting no change in reflectance in this case. For comparisons of average discriminability in scene blocks (Figure 7d), observers were better at discriminating illumination changes than reflectance changes (flat: *t*(8) = 2.42*, p* = 0.046; 3D matte: *t*(8) = 4.41*, p* = 0.007; 3D glossy: *t*(8) = 2.81*, p* = 0.046). Discriminability of reflectance changes was better for the flat scene compared to the 3D matte scene (*t*(8) = 4.23*, p* = 0.009). Discriminability of illumination changes did not show a significant difference between scene blocks.

**Figure 7:**
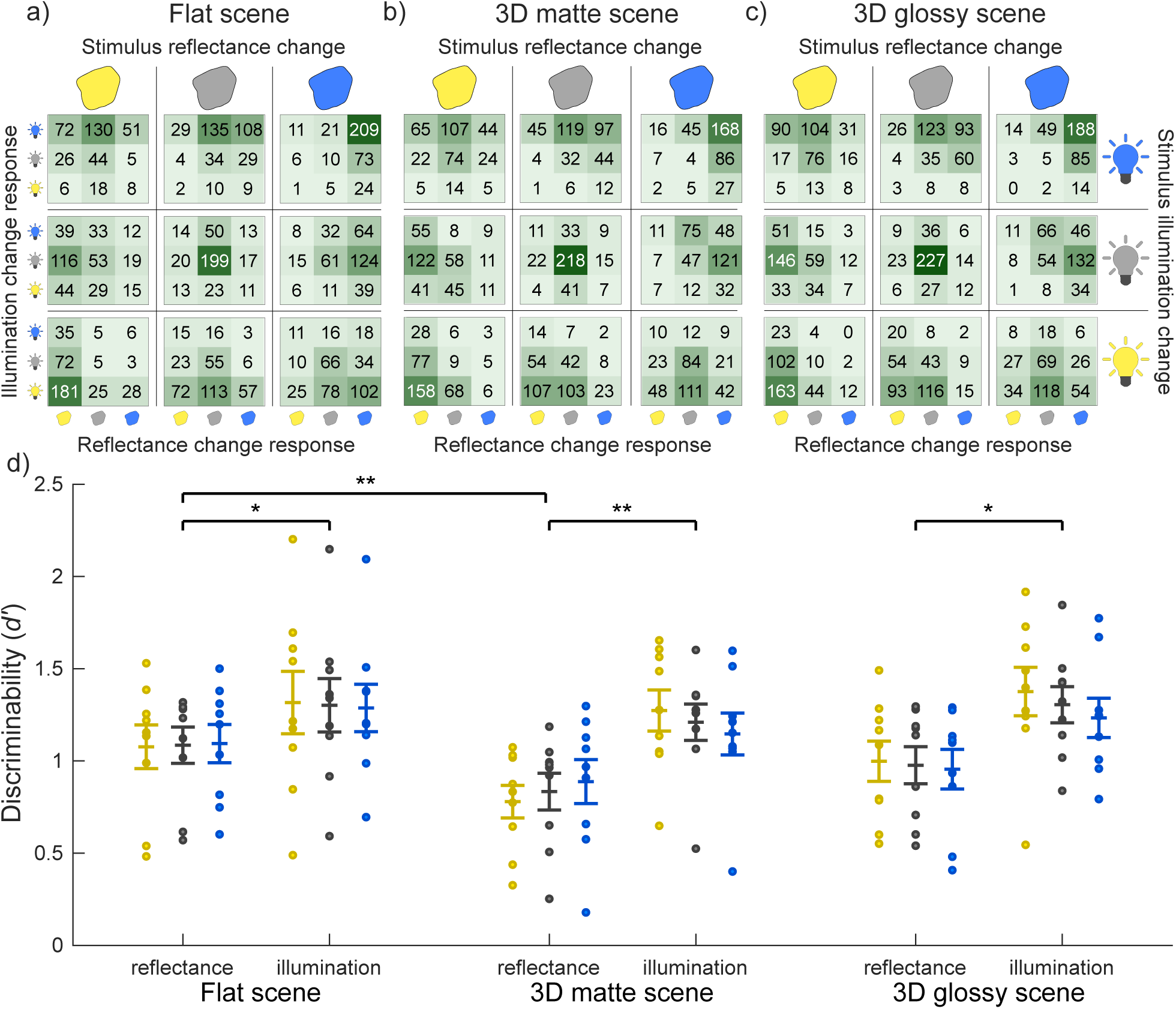
Change identification results from scene blocks. Heatmaps of response counts across all observers for: a) flat scene, b) 3D matte scene, c) 3D glossy scene. Each stimulus change is shown in a separate heatmap. Top icons indicate the correct reflectance change response, and right icons the correct illumination change response. Left icons indicate observers’ illumination responses and bottom icons indicate observers’ reflectance responses. d) plots discriminability measured by average d’ across stimulus reflectance and illumination changes, respectively. Different scene blocks with reflectance and illumination plotted separately are on the x-axis with *d′* on the y-axis. Each dot is a single observer, and the error bars indicate ±1 SEM. Yellow dots show averages across changes towards yellow, blue dots towards blue, and gray the total average. ** *p <* 0.01, * *p <* 0.05.

To illustrate how correct identification of reflectance change depends on the correct identification of illumination change, cross-tabulated responses of scene blocks are presented in Figure 8a–c. Chi-square tests clearly rejected the null hypothesis of independence for the 3D matte (*χ*^2^(3240) = 14.75*, p <* 0.001) and 3D glossy (*χ*^2^(3240) = 35.06*, p <* 0.001) scene blocks, and less decidedly for the flat scene block (*χ*^2^(3240) = 4.41*, p* = 0.036). The independence of correct identifications varied between different color change comparisons: across all scene blocks, independence was not rejected for comparisons where illumination change was accompanied with either a change toward blue or no change in reflectance, but it was rejected for all other comparisons (all *p <* 0.05). Overall, the chi-square tests rejected independence more often than not. However, on average, when the illumination change was not correctly identified, the reflectance change was still correctly identified half of the time — compared to the chance level of one third. Additionally, average conditional *d*′ are presented in Figure 9. In all scenes, discriminability was better with correct identification of illumination change, but only if there was no illumination change in the stimulus (flat: *z* = 3.33*, p <* 0.001; 3D matte: *z* = 3.60*, p <* 0.001; 3D glossy: *z* = 3.85*, p <* 0.001). Otherwise, discriminability did not show a statistically significant difference and performance was always clearly above chance level.

**Figure 8:**
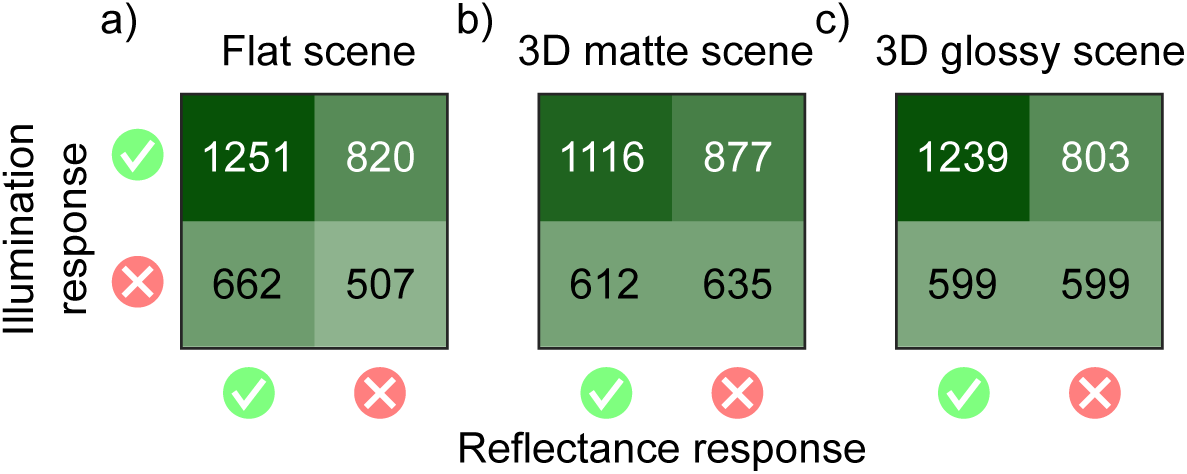
Identification of reflectance changes in scene blocks depending on identification of illumination changes. Response counts across all observers for: a) flat scene, b) 3D matte scene, c) 3D glossy scene, cross-tabulated based on the correctness of the illumination response (row), and reflectance response (column).

**Figure 9:**
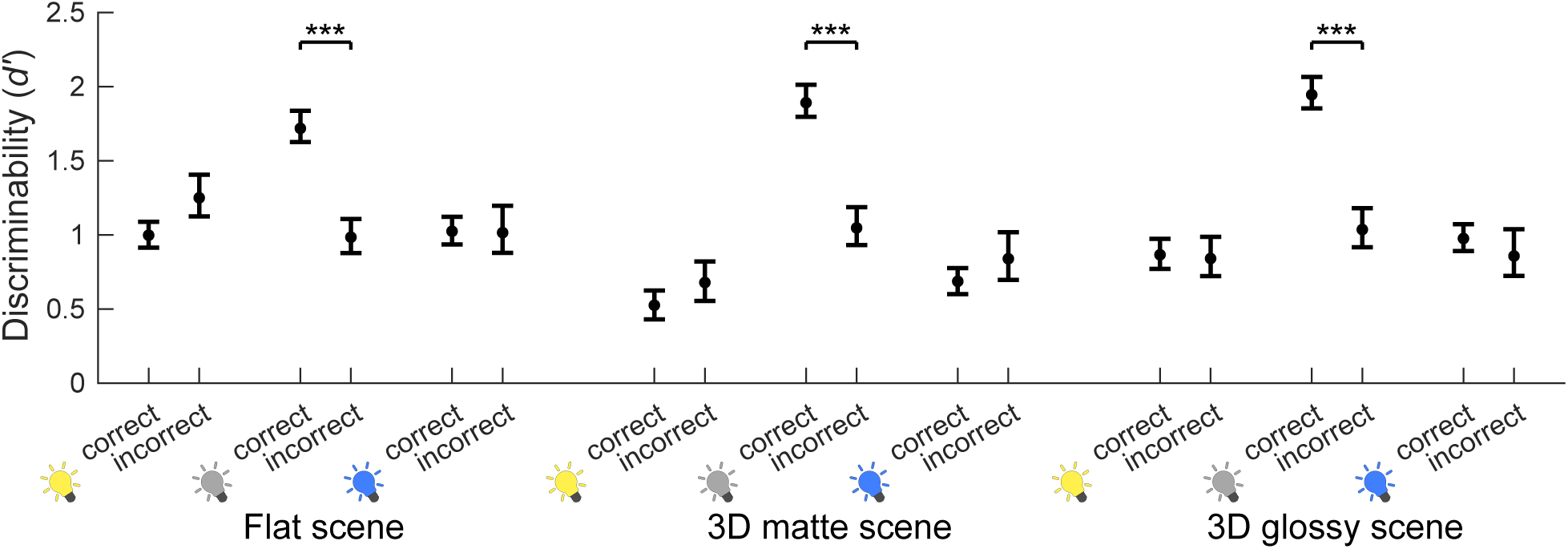
Discriminability of reflectance changes in scene blocks measured by average *d′* in trials with correct and incorrect illumination change identification, respectively. Correct and incorrect illumination change identification for yellow, no, and blue illumination changes within flat, 3D matte, and 3D glossy scene blocks are on the x-axis with *d′* on the y-axis. Error bars display one standard deviation of the mean, calculated by bootstrapping with 10000 iterations and taking the range of the middle 68^th^ percentile. *** *p <* 0.001

Next, we binned the trials based on the reference reflectance and illumination into three categories: the yellow, neutral, and blue categories. We then computed discriminabilities *d*′ and response criteria *c* for reflectance and illumination changes. When trials were split by reference illumination, the decision criterion (Figure 10a) was lower for illumination changes away from neutral than towards neutral (yellow reference: *z* = −12.52*, p <* 0.001; blue reference *z* = 15.80*, p <* 0.001). Note that as all combinations of reference value and stimulus change were analyzed separately, the response criteria for blue and yellow changes are expected to change in opposite directions: for a yellow reference value, a yellow change was away from neutral and a blue change toward neutral, and vice versa for a blue reference. A response criterion of 1 corresponds an optimal or unbiased criterion and a criterion below 1 indicates a bias to respond with the respective color change. The observed criterion shifts thus mean that observers were more likely to respond that the illumination change was away from the neutral color than towards it. In conjunction, discriminability for illumination change (Figure 10c) was higher for changes away from neutral than towards neutral (yellow reference: *z* = −5.35*, p <* 0.001; blue reference *z* = −3.54*, p <* 0.001). In other words, observers were not only more likely to respond that the illumination change was away from neutral, they were also more sensitive to these changes. The response criteria for reflectance (Figure 10b) followed a similar pattern as a function of reference illumination (yellow reference: *z* = −3.40*, p* = 0.001; blue reference *z* = 3.96*, p <* 0.001), but there was no significant difference in sensitivity for reflectance changes (Figure 10d).

**Figure 10:**
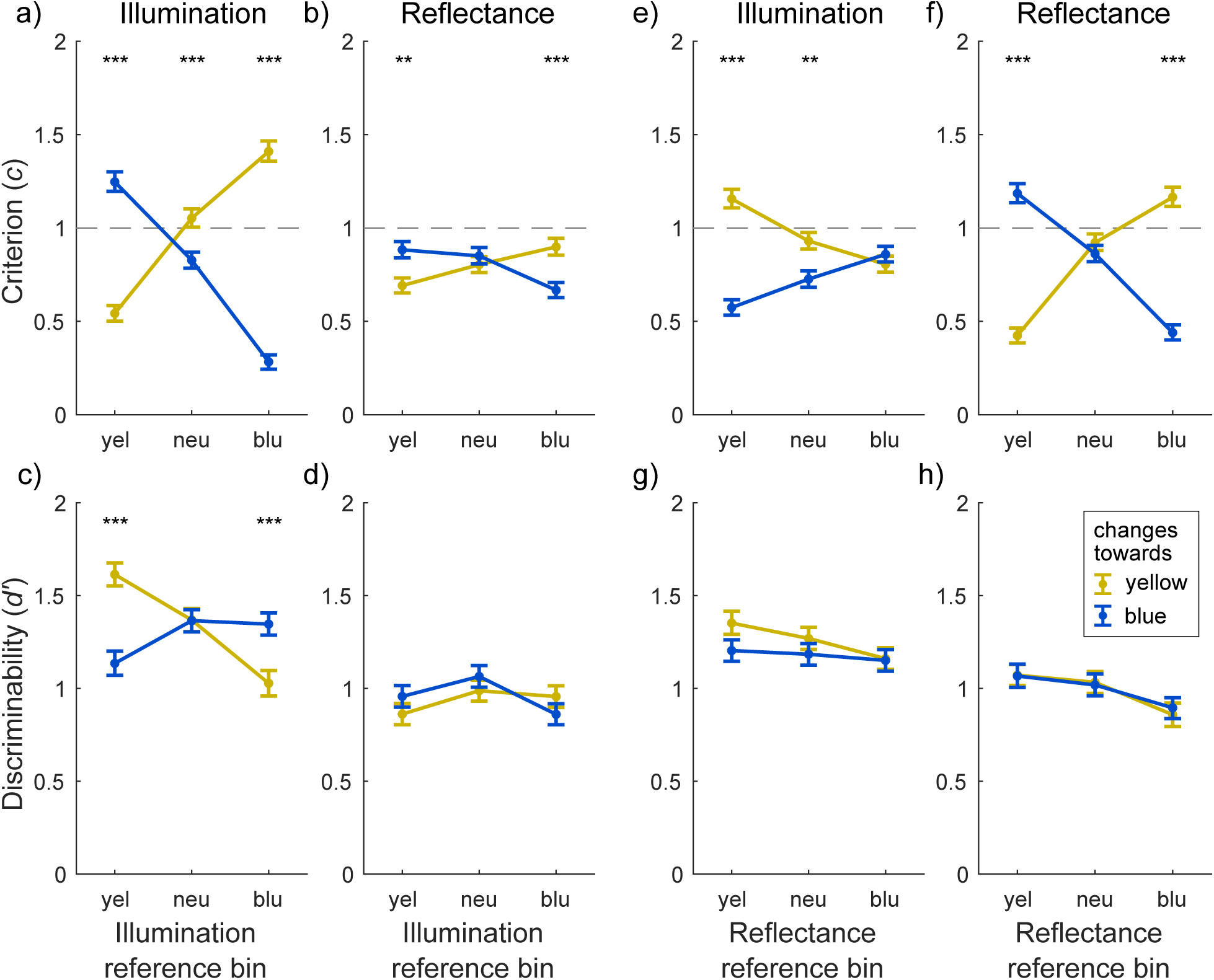
Marginal decision criteria and discrimination indices calculated across all observers for data binned by reference stimulus colors, and reference-comparison color changes. Trials were binned according to whether the reference had illumination (a–d), or reflectance (e–h), from the yellowest, bluest, or neutral third in the stimulus pool (x-axis). a–b) and e–f) show criterion (*c*, y-axis), and c–d) and g–h) show discriminability (*d*’, y-axis), for identification of illumination and reflectance changes, respectively. *d*’ and *c* were calculated between identification of no change, and yellow or blue change (yellow and blue lines). Error bars display one standard deviation of the mean, calculated by bootstrapping with 10000 iterations and taking the range of the middle 68^th^ percentile. Note that criteria are displayed in absolute values and a criterion below 1 means a bias to respond with the change indicated by the line color. *** *p <* 0.001, ** *p <* 0.01.

When trials were binned by reference reflectance, the criteria for reflectance (Figure 10f) were higher for changes towards neutral than away from neutral (yellow reference: *z* = −13.96*, p <* 0.001; blue reference *z* = 11.44*, p <* 0.001). For illumination changes, the pattern was different (Figure 10e): the criterion was higher for illumination changes towards yellow in the yellow reference group (*z* = 9.11*, p <* 0.001). The observers were thus more likely to report blue illumination changes when the reference reflectance was yellow. There were no significant differences in sensitivity for either reflectance or illumination changes (Figure 10g–h).

Finally, we quantified the amount of cross-talk in the processing of reflectance and illumination cues by fitting a two-dimensional signal-detection model to the data. The model has two hypothetical mechanisms, a “reflectance mechanism” and an “illumination mechanism”, whose directions in stimulus space determine how the two cues affect the responses. The raw data for the model fits were the response counts from the scene blocks, shown in Figure 7a–c. Note that this model does not take into account the asymmetries demonstrated in Figure 10, nor does it predict the specific pattern of conditional *d*′-values in Figure 9. The vectors depicting the mechanisms in stimulus space are shown in Figure 11a for the three scene conditions. The axes represent the two possible stimulus changes, a reflectance change and an illumination change, with a blue change as positive and a yellow change as negative. If there was no cross-talk in the identification of the two types of change, the vectors would lie on the corresponding axes.

**Figure 11:**
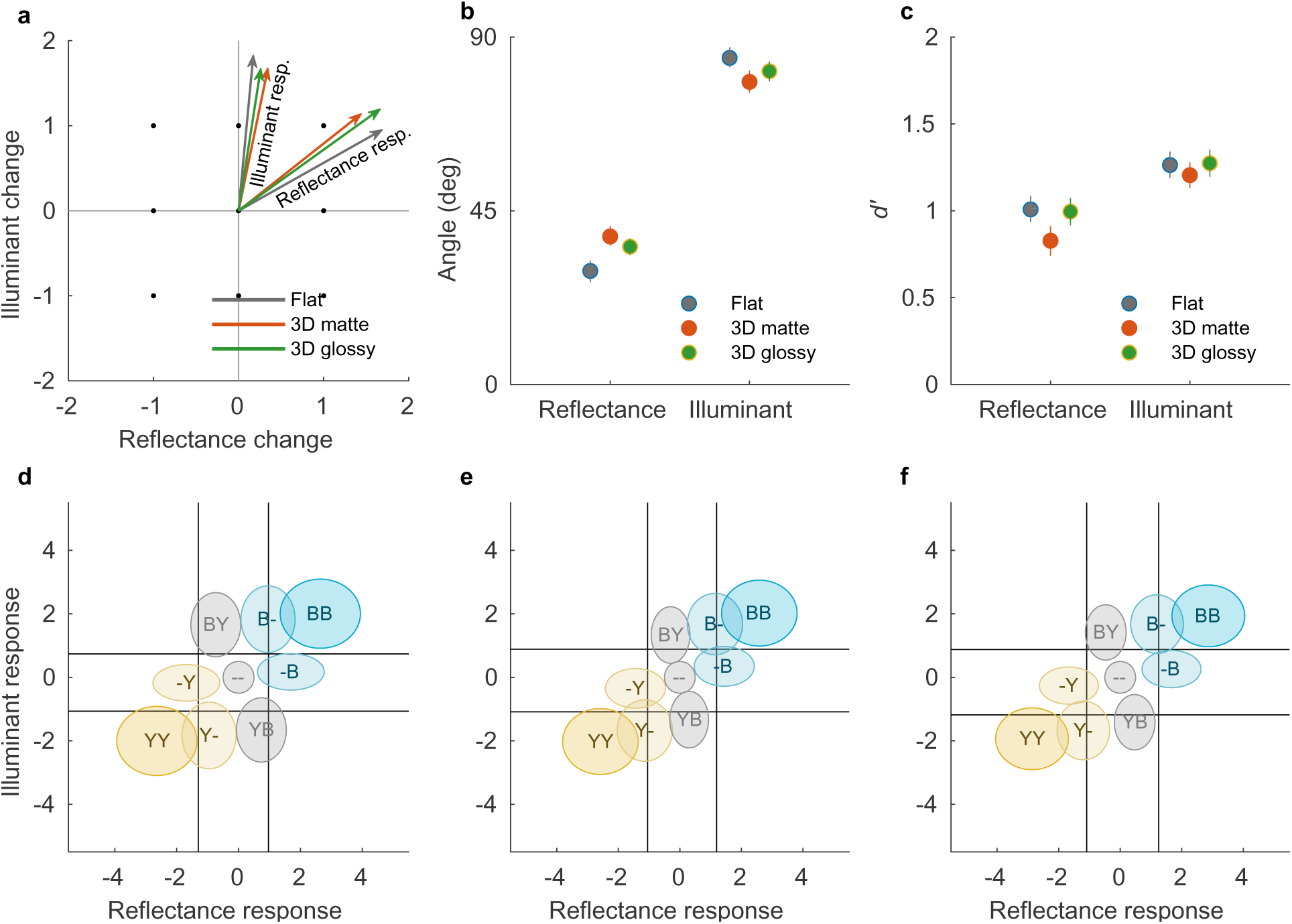
a) The vectors depicting the reflectance and illumination mechanisms in stimulus space for the three scene conditions: flat, 3D matte, and 3D glossy. The more aligned a given vector is with the corresponding stimulus axis, the better the model observer is at identifying the changes, and the fewer confusions between the two types of stimulus change occur. Dots indicate the nine stimulus changes used in the experiment. b) The angle of the mechanism vectors with respect to the reflectance axis. c) Average *d′*-values of the model observer for identifying reflectance and illumination changes. d–f) Response distributions for the nine types of stimulus change plotted in the decision space of the model (the axes correspond to the mechanism responses) in the flat (d), 3D matte (e) and 3D glossy (f) conditions. The vertical and horizontal lines show the response criteria. Ellipses signify half a standard deviation of the response distribution around the mean. First letter in each distribution indicates stimulus illumination change, second letter reflectance change. Color of the circle indicates absolute color change of the target.

Instead, in all three conditions, neither mechanism vector is aligned with the axis. This means that illumination cues caused a response in the reflectance mechanism, and vice versa. The illumination mechanism was, however, much more closely aligned with the illumination change axis in the stimulus space. In other words, the reflectance mechanism was more excited by illumination changes than the illumination mechanism was by reflectance changes. Thus, the illumination responses are much more independent of reflectance color changes than the other way around. In the 3D matte scene, the reflectance mechanism is close to having a 45^◦^ angle with the axes, which would indicate that both cues were equally likely to result in a perceived reflectance change. We tested the separation of the reflectance and illumination mechanisms with permutation tests. Compared to the 3D matte scene, the angle between the two mechanisms was larger in the presence of local contrast (flat scene, *p <* 0.01) or specular highlights (3D glossy scene, *p <* 0.05), leading to better separation of the two dimensions. The angles between the mechanism vectors and the reflectance axis (x-axis) are plotted in Figure 11b. The average *d*′-values are plotted in Figure 11c. Although grouped differently, these correspond to the average *d*′-values in Figure 7d and show a similar pattern, with poorest color constancy for the 3D matte stimulus.

Figure 11d–f show the response distributions for each stimulus change in decision space. The decision criteria are shown as vertical and horizontal lines. These distributions further illustrate how reflectance responses are much more dependent on illumination changes than illumination responses are on reflectance changes.

## 3 Discussion

We examined whether operational color constancy relies on explicit identification of illumination changes, and the contribution of different cues to color constancy. Comparing average discriminability in scene blocks (Figure 7d), we found that illumination changes were more reliably discriminated than reflectance changes, and that local contrast improved color constancy of reflectance percepts. Discriminability suggested that specular highlights might also produce a small improvement in color constancy and this was seen in better separation of modeled mechanism vectors (Figure 11). When we examined discriminability of reflectance changes depending on correct identification of illumination changes (Figure 9), we found that identification of a reflectance change was improved if observers correctly reported an illumination change only when there was no illumination change in the stimulus. When there was an illumination change in the stimulus, discriminability of reflectance changes was independent of correct identification of the illumination change. By comparing discriminability of color changes for different reference colors (Figure 10), we found that there was a bias to more readily report changes away from the neutral point, both with illuminations and reflectances, and also better discriminability of illumination changes away from the neutral point. Broadly, local contrast and specular highlights seem to improve color constancy, that is, they help to disambiguate illumination and surface reflectance. However, they do so without providing any better explicit judgment of illumination chromaticity.

In our experiment, identification of illumination changes was more accurate than identification of reflectance changes (Figure 7d), which is in contrast with previous studies finding poorer judgments of illumination than reflectance chromaticity (Granzier et al., 2009; Rutherford & Brainard, 2002). This is probably caused by the experimental design; illumination change could be identified directly from a global chromaticity shift alone (ignoring the target) while identification of reflectance change required comparison between target and context. In contrast to our findings, in the study by Ennis and Doerschner (2019), observers’ responses of color change followed real stimulus changes approximately equally well for illumination and reflectance. This is likely due to the type of chromaticity changes employed. As we chose illumination and reflectance chromaticity changes that would lead to the same reflected chromaticities, the step sizes for both changes had to be identical. Instead, Ennis and Doerschner (2019) used significantly larger chromaticity changes for reflectance than for illumination. The changes were also not exactly matched for hue and included combinations of illumination and reflectance changes along different axes (red-green, blue-yellow). We deliberately chose illumination and reflectance chromaticity changes that can cancel each other out in the light reflected from the target object to study the dependence of operational color constancy on illumination change judgments. Correct reflectance change judgments are in theory conditional on correct illumination change judgments by the experiment design. However, unlike in previous studies which measured the accuracy of illumination chromaticity estimates (Granzier et al., 2009) or constancy by illumination chromaticity adjustments (Rutherford & Brainard, 2002), here the task only required identifying if a possible change in illumination was towards yellow or blue. Despite these design choices, we found that in most cases discriminability of reflectance changes did not depend on correctly identifying the illumination change (Figure 9). Irrespective of the stimulus scene, only trials with no illumination change showed a significant dependence. It is not clear what causes this pattern of results. In the task, observers could identify the illumination change directly from the context, without having to consider the target at all. As such, if there was no illumination change, an incorrect response would either require that a change in the target only is misattributed as a change across the whole stimulus, or a momentary lapse in performing the task. It may be that misattributing reflectance changes as illumination changes represents a more fundamental failure in parsing chromatic changes in the scene, leading to an accumulation of trials where performance was overall poor. Regardless, discriminability of reflectance changes clearly did not fully depend on illumination change judgments.

The illumination-estimation hypothesis posits that the visual system estimates scene illumination to determine surface color (see, e.g., Maloney & Yang, 2003) and errors in reflectance estimates could be a consequence of errors in illumination estimates (Brainard, 1998; Brainard et al., 1997). Studies characterizing explicit illumination judgments, however, do not fully support this view (Granzier et al., 2009; Rutherford & Brainard, 2002; Weiss et al., 2017). It is still plausible that the full computation to estimate illumination is implicit and as the visual system attempts to effectively discount scene illumination, perceived illumination chromaticity is consistently desaturated (as seen in Granzier et al., 2009). However, our results suggest that color constancy is mostly disconnected from even a rough explicit illumination estimate in the form of identifying a possible illumination change towards yellow or blue.

To examine the role of local contrast and specular highlights in disentangling illumination and reflectance changes, these cues were varied in scene blocks by manipulating the scene background. The flat scene included a background behind the target, creating local contrast at the object outline. Instead, with the 3D matte scene, illumination changes had to be interpreted across a larger spatial distance. The lack of local contrast was produced by having the objects, as explained to observers: “float in a black void.” Note that this is different from studies that manipulated local contrast cues by introducing a conflicting cue instead (Delahunt & Brainard, 2004; Kraft et al., 2002; Xiao et al., 2012). An invalid local contrast cue does not only eliminate the cue, but is likely to interfere with other cues. In the flat scene, the uniformly gray background reflected illumination chromaticity unaltered. In contrast to the 3D matte scene in which illumination changes had to be interpreted from an average chromaticity change in the scene, the glossy scene included specular highlights that could be used as a direct cue to illumination chromaticity similarly to the flat scene. However, specular highlights had to be spotted from specific locations across larger spatial distances. These manipulations do not provide a full factorial breakdown of the visual background cues in our stimuli, but are aimed towards two comparisons specifically: local contrast vs. no local contrast, and accurate illumination chromaticity cue vs. averaging.

Our results show that discriminability of illumination changes did not differ between the scene blocks (Figure 7d). This is somewhat surprising. Color constancy is improved with the inclusion of visual cues such as local contrast (Delahunt & Brainard, 2004; Kraft & Brainard, 1999; Kraft et al., 2002) and specular highlights (Granzier et al., 2014; Lee & Smithson, 2016; Wedge-Roberts et al., 2020). The visual cues can be thought to provide the visual system with more information of scene illumination (e.g., Boyaci et al., 2006; Yang & Maloney, 2001) which the visual system may further combine for more accurate and reliable illumination estimates (Maloney, 2002). Yet we did not find a significant benefit in discrimination of illumination changes with either local contrast or specular highlights. Again, explicit identification of illumination changes even with more reliable cues seems limited.

What differed between the scene blocks was the discriminability of reflectance changes, being highest for the flat scene and lowest for the 3D matte scene (Figure 7d). Previous studies have demonstrated the importance of local contrast for accurate color constancy (Delahunt & Brainard, 2004; Kraft & Brainard, 1999; Kraft et al., 2002), but by introducing a conflicting cue they produced a dramatic deterioration in constancy. Instead, we found only a medium–large benefit with inclusion of local contrast (reflectance discriminability between flat scene and 3D matte scene, Cohen’s *d* = 0.76). Glossiness, or specular highlights, also supports color constancy, especially when cues are limited (Granzier et al., 2014; Lee & Smithson, 2016; Wedge-Roberts et al., 2020). For specular highlights, we only found a small–medium benefit (reflectance discriminability between 3D glossy scene and 3D matte scene, Cohen’s *d* = 0.43, not statistically significant). These findings suggest that the visual system does not need to rely on local contrast or direct illumination chromaticity cues, but can achieve a decent degree of color constancy even by averaging and/or comparison across spatially disjointed elements.

The observed data was also well accounted for by modeling hypothetical mechanisms for illumination and reflectance response. Note that these hypothetical mechanisms were defined specifically for the stimulus and response space in this experiment design. They only abstract the association between stimulus and response with no assumptions of underlying sensory or cognitive mechanisms. In contrast to studies of operational color constancy where the choice is between either a change in reflectance or illumination (Craven & Foster, 1992), or between congruent and incongruent changes (Foster, Nascimento, et al., 2001; Linnell & Foster, 1996), simultaneous identification (also in Ennis & Doerschner, 2019) introduces different requirements for the visual system. A uniform chromaticity shift in the stimulus can still signal that there was an illumination change without reflectance change. However, a non-uniform chromaticity shift can signal a reflectance change, but it can occur with or without an accompanying illumination change. Furthermore, the target reflectance can change similarly or inversely compared to illumination, producing either a large shift or no shift in the target, respectively. Neither is uniquely defined by a global or local (target) chromaticity shift alone. Responses that fail to separate changes in illumination and reflectance from this ambiguity lead to cross-talk between the model mechanisms.

Based on model estimates (Figure 11), in all scene blocks, the illumination response was well aligned with illumination changes in the stimulus, indicating that reflectance changes were rarely misidentified as illumination changes. In contrast, the reflectance response was much more mixed between reflectance and illumination changes, indicating that changes in illumination were often misidentified as changes in reflectance. The independence of the reflectance mechanism in the modeled vectors mirrors the difference in reflectance identification we found between the scene blocks. However, they also show a difference for the illumination mechanism between the scene blocks while difference in illumination identification *d*′ did not reach statistical significance. Thus, even if local contrast and specular highlights do not produce a notable advantage for judging illumination changes in operational color constancy, they seem to help the visual system resolve at least some ambiguity. This further supports the notion that the visual cues tested here support the disentangling of illumination and reflectance, not absolute illumination chromaticity judgments.

On average, there was no difference in discriminability for changes towards yellow and changes towards blue. However, this only held when averaged over all reference chromaticities. When different illumination reference chromaticities were analyzed separately, discriminability was higher and the decision criteria lower for illumination changes away from neutral compared to changes towards neutral (Figure 10). This bias in discriminability is in agreement with the “neutral bias” coined by Hurlbert and colleagues (Pastilha et al., 2020; see also Aston et al., 2019), but our results also demonstrated a strong complementary bias in the decision criteria, which was not reported in previous studies. Furthermore, the bias in decision criteria was not limited to illumination changes but was also found for reflectance changes towards and away from neutral depending on reflectance reference. Thus, there seems to be a significant component to the neutral bias driven by decision criteria which is not limited to percepts of illumination chromaticity, but influences surface color percepts as well.

As discussed by previous studies that identified a neutral bias in illumination discrimination (Aston et al., 2019; Pastilha et al., 2020), the bias is not easily explained by low-level asymmetries of chromatic discrimination. For example, while detection thresholds for changes along the L - M cone-opponent axis are independent of the adaptation point on this axis, detection thresholds on the S - (L + M) axis increase with increasing S-cone excitation of the adapting stimulus (Krauskopf & Gegenfurtner, 1992). However, this means that changes towards both yellow and blue are easier to detect against a yellow background, whereas in the neutral bias, changed towards yellow are easier to detect from a yellow reference, and changes towards blue are easier to detect from a blue reference. Another plausible explanation is a strong prior for illumination chromaticity in a Bayesian framework of color constancy (Brainard & Freeman, 1997; Brainard et al., 2006; Gehler et al., 2008), as was suggested by Hurlbert (2019). The range of illumination chromaticity in our experiment was along the common axis of natural illumination variation of the daylight locus (Judd et al., 1964) and by extension natural scene color variation (McDermott & Webster, 2012; Webster & Mollon, 1997; Webster et al., 2007). With respect to natural variation, however, our "neutral" references are still much more common than the saturated "yellower" and "bluer" references. As memory retention can increase the weighting of priors in color estimates (Olkkonen & Allred, 2014; Olkkonen et al., 2014), this would produce a stronger influence of a neutral prior in the reference stimulus than the comparison stimulus. This would then lead to both a lower criterion and higher sensitivity for identifying changes away from neutral. But if this prior is specific to illumination chromaticity, why do we also see a bias in the reflectance criterion? A speculative answer is that the bias for illumination judgments is partially transferred to reflectance judgments. We found that there is a strong interference of illumination changes in the stimulus on reflectance responses (Figure 11), but it is unclear if this could also influence the bias that depends on reflectance reference color (Figure 10f). Regardless, to confirm if an illumination prior could underlie the pronounced neutral bias in illumination estimation requires further study.

## 4 Conclusion

Color constancy does not fully depend on correctly identifying the illumination chromaticity. Furthermore, cues from local contrast and specular highlights aid color constancy in accurate disentanglement of reflectance from illumination, but not through improved illumination chromaticity judgments. Finally, changes away from the chromatic neutral point are not only better discriminated for illuminations, but also more readily reported for both illuminations and surface colors.

## Acknowledgements

This project was funded by Emil Aaltonen Foundation (Grants #200240 N1 and #210249 N1).

